# High-Dimensional Representation of Texture in the Somatosensory Cortex of Primates

**DOI:** 10.1101/451187

**Authors:** Justin D. Lieber, Sliman J. Bensmaia

## Abstract

In the somatosensory nerves, the tactile perception of texture is driven by spatial and temporal patterns of activation distributed across three populations of afferents. These disparate streams of information must then be integrated centrally to achieve a unified percept of texture. To investigate the representation of texture in somatosensory cortex, we scanned a wide range of natural textures across the fingertips of Rhesus macaques and recorded the responses evoked in Brodmann’s areas 3b, 1, and 2. We found that texture identity is reliably encoded in the idiosyncratic responses of populations of cortical neurons, giving rise to a high-dimensional representation of texture. Cortical neurons fall along a continuum in their sensitivity to fine vs. coarse texture, and neurons at the extrema of this continuum seem to receive their major input from different afferent populations. Finally, we show that cortical responses can account for several aspects of texture perception in humans.

## Introduction

Our sense of touch endows us with an exquisite sensitivity to surface microstructure. We can perceive surface features that range in size from tens of nanometers (Skedung et al., 2013) to tens of millimeters and integrate these to form a cohesive textural percept. In the somatosensory nerves, surface features at different spatial scales are encoded in different populations of afferents and rely on different neural representations. Coarse surface features are reflected in the spatial patterns of activation evoked in slowly adapting type-1 (SA1) and rapidly adapting (RA) afferents, whose small receptive fields give rise to a faithful neural image of surface elements measured in millimeters (Johnson and Lamb, 1981; Phillips and Johnson, 1981). However, many tangible surface features are too small and too close together to be encoded spatially because the spatial code is limited by the innervation density of the skin (Darian-Smith and Kenins, 1980; Johansson and Vallbo, 1979). To perceive fine textural features requires movement between skin and surface, which leads to the elicitation of texture-specific skin vibrations, which in turn evoke precisely timed texture-specific spiking patterns in RA and Pacinian corpuscle-associated (PC) afferents (Bensmaïa and Hollins, 2005; Hollins and Risner, 2000; Hollins et al., 2001, 2002; Lieber et al., 2017; Weber et al., 2013). These spatial and temporal representations must be combined and synthesized to achieve a unified percept of texture, a process about which little is known.

While neurons in somatosensory cortex have been shown to encode information about texture, previous studies investigating cortical texture representations used surfaces with elements in the range of millimeters, such as Braille-like dot patterns (DiCarlo and Johnson, 2000; DiCarlo et al., 1998) and gratings (Darian-Smith et al., 1982; Sinclair and Burton, 1991; Tremblay et al., 1996), which span only a small fraction of the wide range of tangible textures. We have previously shown that responses to such textures – which only engage the spatial mechanism – provide an incomplete view of the neural mechanisms that mediate the perception of texture (Lieber et al., 2017; Weber et al., 2013).

To fill this gap, we examined how textures that span the tangible range are encoded in somatosensory cortex. To this end, we scanned a wide range of textures – including fabrics, furs, and papers, in addition to the traditional embossed dots and gratings – across the fingertips of (awake) Rhesus macaques and recorded the responses evoked in somatosensory cortex, including Brodmann’s areas 3b, 1, and 2. First, we found that texture identity is faithfully encoded by these neuronal populations and that texture information is distributed across neurons which each exhibit idiosyncratic texture responses. Second, we showed that the heterogeneity across somatosensory neurons is in part driven by differences in the submodality composition of their input (SA1, RA, PC). We then discovered the downstream recipients of the spatial and temporal codes observed at the periphery: a subpopulation of cortical neurons receives strong input from SA1 fibers and preferentially encodes coarse textural features whereas another population of neurons receives strong input from PC fibers and preferentially encodes fine surface features. Finally, we showed that the responses of somatosensory neurons account for psychophysical reports of texture obtained from human observers.

## Results

We recorded the activity evoked in 141 neurons in somatosensory cortex (35 from area 3b, 81 from area 1, and 25 from area 2) from three Rhesus macaques with receptive fields on the distal fingertip, as each of 59 textured surfaces (Table S1) was scanned across the skin using a rotating drum stimulator, which allows for precise control of scanning speed and indentation depth (Figure 1A-B). These surfaces were chosen to vary widely in microstructure and material properties in an attempt to explore as fully as possible the range of everyday textures. The objective of the study was to determine the degree to which texture information is encoded in cortex, examine the nature of this representation, and assess the degree to which this representation can account for perception.

**Figure 1.**
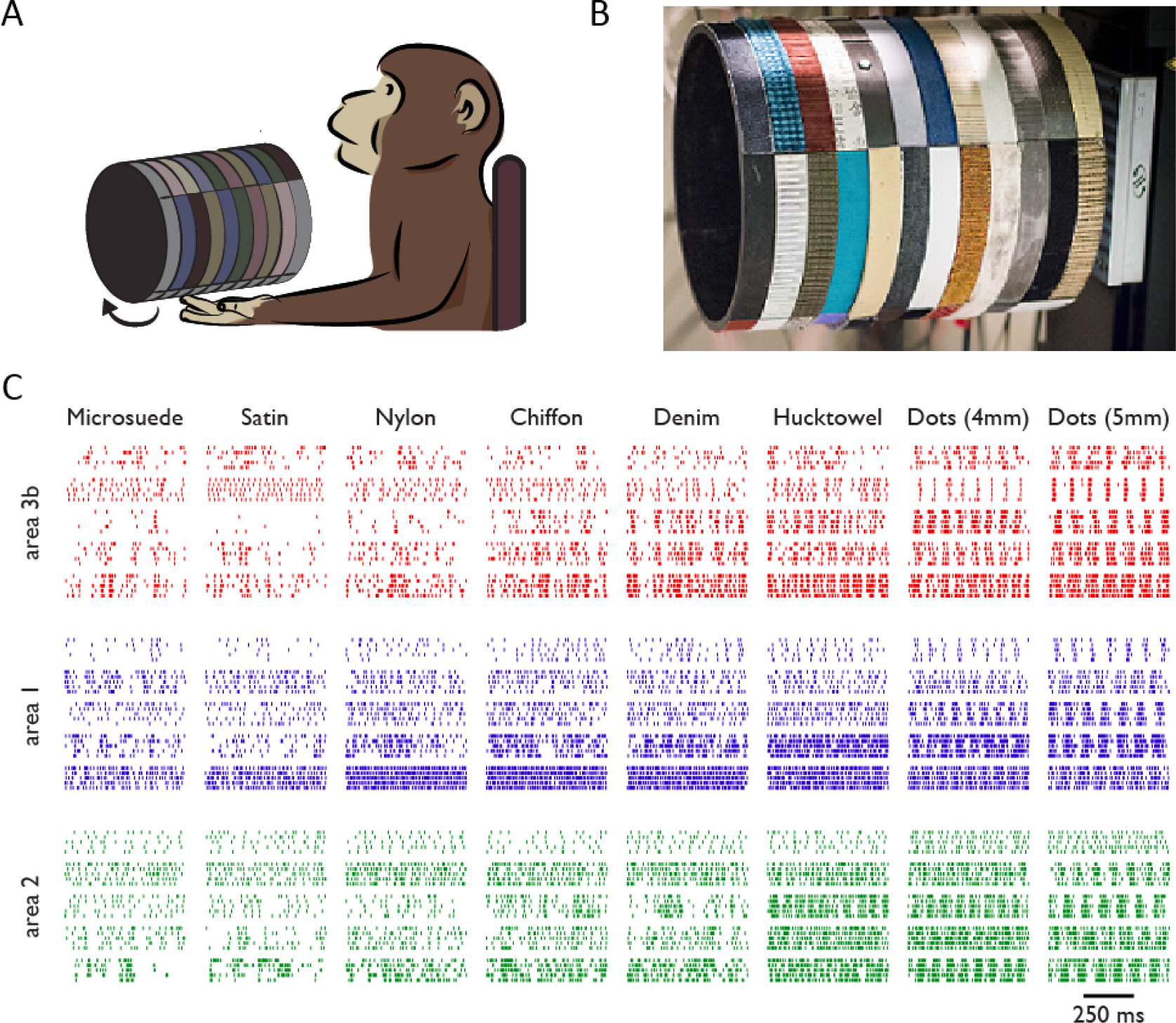
Experimental apparatus and sample texture responses. A| Textures were passively presented to the distal fingerpads of awake macaques. B| The rotating drum stimulator – on which the 59 textures were mounted – allows a surface to be scanned across the fingertip at a precise and repeatable speed and depth of indentation into the skin. C| Sample spiking responses of 5 neurons each in areas 3b, 1, and 2 to 5 repetitions of 8 textured surfaces.

### Neurons in somatosensory cortex encode texture

First, we examined the degree to which the responses of individual somatosensory neurons are modulated by texture (Figure 1C). We found that nearly every neuron responded to at least one texture (140/141 neurons modulated above baseline firing rate, *p*<0.05, permutation test with Bonferroni correction), and that each texture significantly modulated the response of at least 20% of the neurons (0.23-0.72-92, min-median-max proportion of textures across cells, permutation test). To test whether these neurons carry texture-specific information, we built a simple linear classifier based on single trial spike counts. Nearly all neurons yielded classification performance that was significantly above chance (mean ± s.d. of performance: 6.7% ±3.7%, chance performance: 1.7%, 95% of neurons > chance), and neurons that yielded better than chance performance were approximately equally prevalent in areas 3b, 1, and 2 (97%, 96% and 88%, respectively, Figure S1A).

Next, we examined the degree to which texture identity is encoded in the responses of populations of somatosensory neurons (Figure 2A). To this end, we implemented the texture classifier using the responses of groups of neurons of varying size. We found that high classification performance could be achieved with a small population of somatosensory neurons (as few as 83 neurons yielded 97% performance) and that the full population yielded nearly perfect performance (Figure 2B). Classification performance was largely comparable across cortical modules (an average of 73%, 72%, 62%, for groups of 25 neurons in areas 3b, 1, 2, respectively, see Figure S1B) and was robust to (simulated) noise correlations (Figure S2A). In summary, small populations of somatosensory neurons convey sufficient information to support texture identification for a large and diverse texture set.

**Figure 2.**
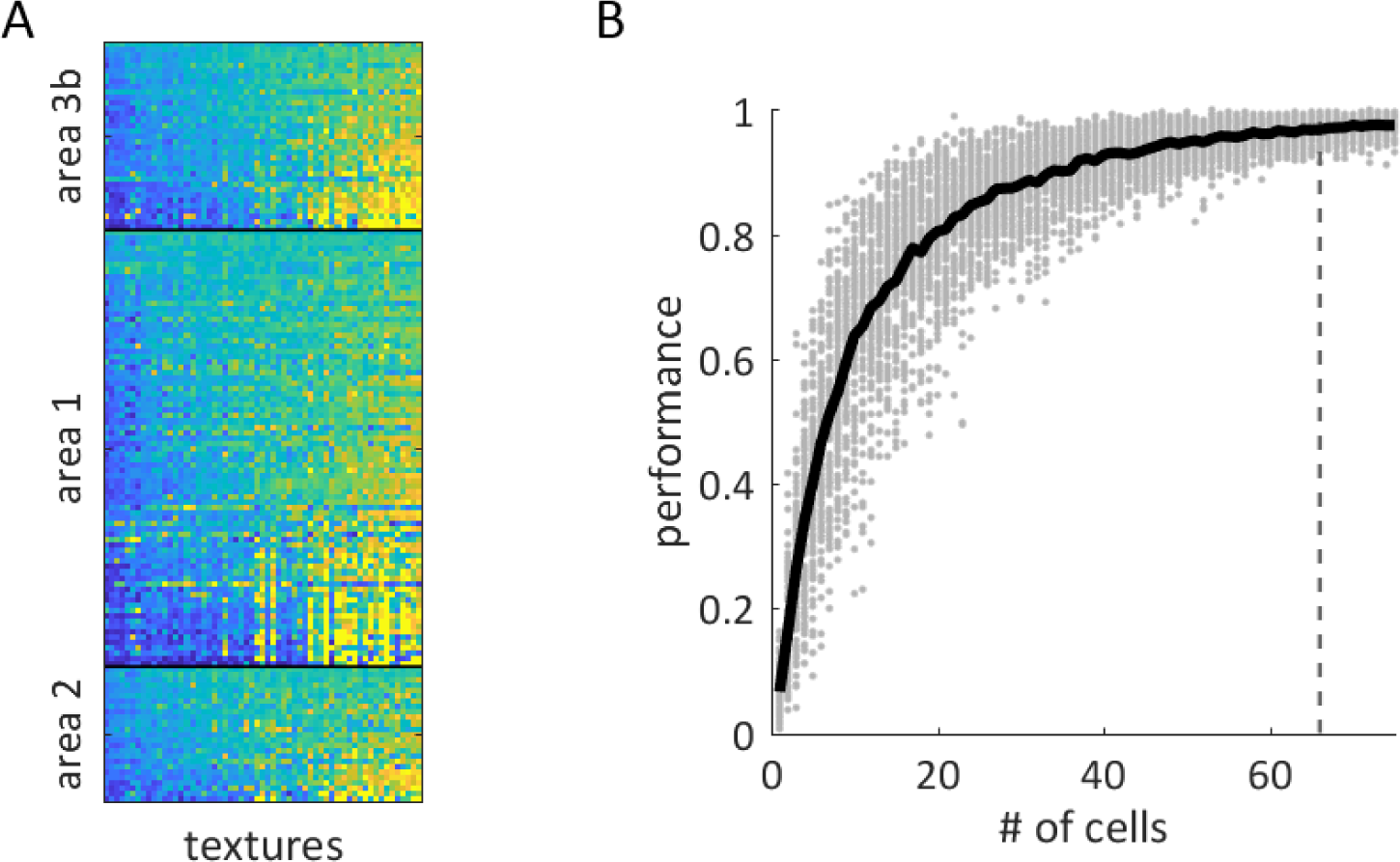
Neurons in somatosensory cortex encode texture. A| The mean firing rate (across repetitions) evoked by each of 59 textures (columns) in somatosensory neurons, split by cortical field. Firing rates are normalized within neuron for display purposes, ranging from low (blue) to high (yellow). Textures are sorted according to the first principal component of the population response from lowest to highest. Cells are ordered first by area, then by variance of their firing rates across textures. Somatosensory neurons exhibit heterogeneous responses to textures. B| Texture classification performance of groups of cortical neurons is plotted against the size of the neuron group. As expected, classification performance improves as more cells were included. Gray dots denote the performance of individual neuronal groups, the black trace denotes the mean as a function of group size. Groups of 66 cells, marked by the grey dashed line, yield a near-asymptotic performance of 97%.

### The cortical representation of texture is high-dimensional

Two factors drive the ability of neural populations to classify stimuli more accurately than do individual cells. First, as more neurons are included, the trial-to-trial variation in response is averaged out. Second, increasing the variety in tuning properties in the neural population can more effectively represent the high-dimensional character of a complex stimulus, and thus increase the effective dimensionality of the resulting neural representation. That is, insofar as different neurons respond to different aspects of a surface, these idiosyncratic responses will provide information beyond that available from simply averaging responses across cells.

We examined the dimensionality of texture responses – the degree to which somatosensory neurons respond heterogeneously to texture – by performing a principal components analysis (PCA) on the population response. That is, we first characterized the correlational structure in texture responses across neurons in somatosensory cortex and then assessed the degree to which responses could be reduced to a smaller set of non-redundant signals. We found that most of the variance in neuronal responses was explained by the first principal component (Figure 3A) (65% proportion of variance explained from the 1st component, essentially the mean population firing rate, *R^2^*=0.99), a signal that was strongly preserved across all three cortical areas (with inter-correlations of first principal components across pairs of areas all greater than 0.95, see Figure S1C). As discussed below, this prominent neuronal dimension has a clear perceptual correlate.

**Figure 3.**
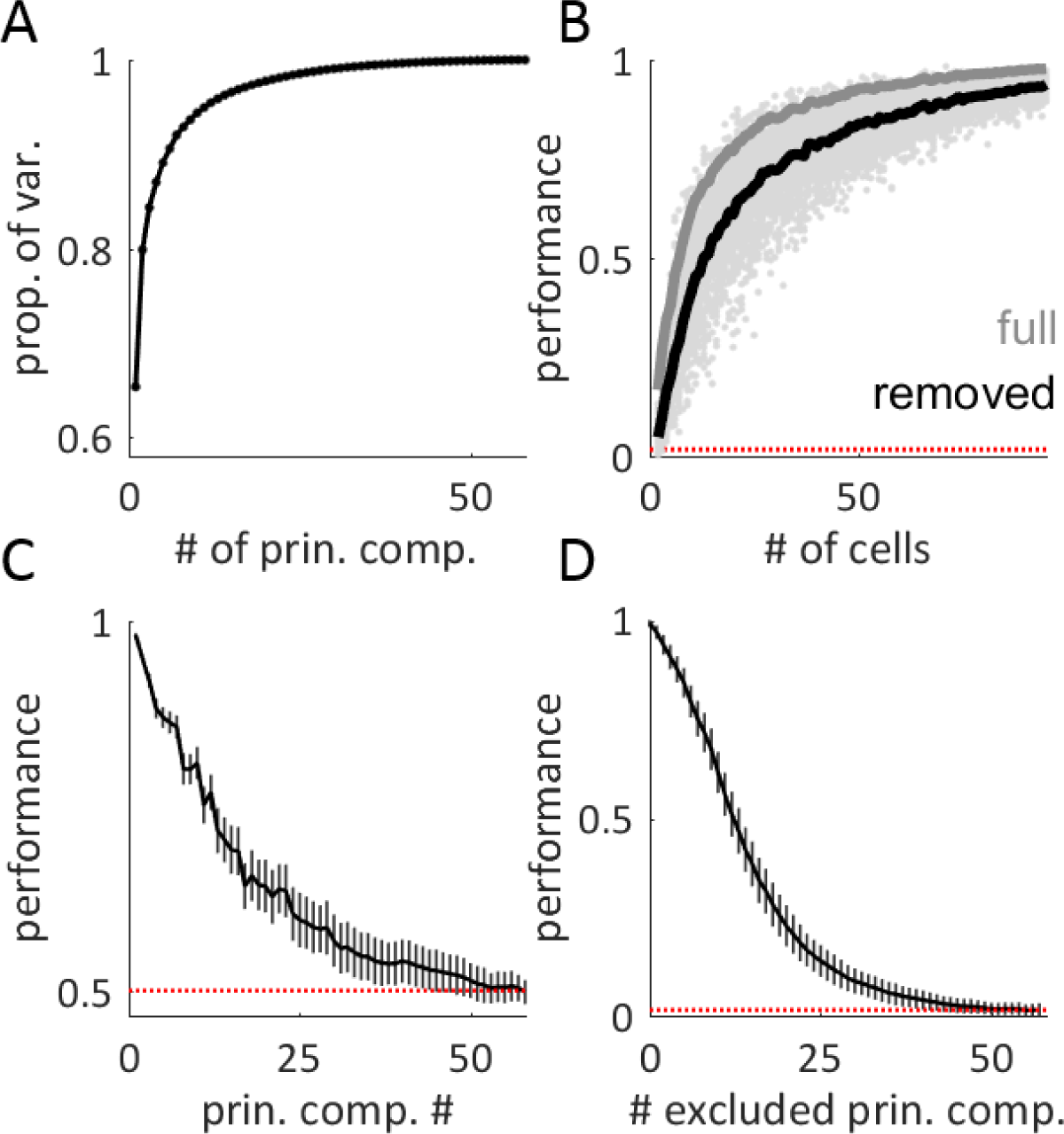
The cortical representation of texture is high-dimensional. A| Cumulative scree plot (proportion of variance explained) for the principal components analysis on the population response to texture. The bulk of the response variance is carried by the first few components. B| Texture classification (as in Figure 2) using the cortical population response with the full population response (grey line) and with the first principal component removed (black line). The red dotted line denotes chance performance. Classification performance is only slightly reduced when this first component is removed. C| Average classification performance of each individual principal component when distinguishing between pairs of textures. Responses were above chance even for components that explained only a small proportion of the total variance. Error bars represent the standard deviation of performance across texture pairs and shuffling of the training and test sets. D| Classification performance based on firing rates projected onto a subset of principal components, built by excluding the N principal components in decreasing order of their eigenvalues (i.e., removing the largest components first). Even when dozens of the high-variance principal components were removed from the response, texture classification was still above chance.

We sought to characterize whether heterogeneity in texture responses across neurons provides texture-specific information beyond that found in the mean population response. To this end, we again implemented the texture classifier, this time using only a subset of the principal components of the neural response. When the population response was collapsed onto a single dimension – the first principal component – classification performance dropped to 41%, compared to 99.4% when the entire response was used. Conversely, if we removed only the first principal component from the population response and preserved all other components, we achieved 92% classification accuracy with as few as 83 cells and 97% accuracy with the full population of 141 cells (Figure 3B). In other words, the heterogeneity of neural responses to texture is a major contributor to the texture signal in cortex.

Given the importance of low-variance dimensions to classification performance, we sought to assess how many of these dimensions are reliably informative about texture identity. To this end, we first quantified how many dimensions identified through PCA reliably carried texture information. We found that the first 30 dimensions carried sufficient texture information to distinguish pairs of textures (Figure 3C, 95% of trial shuffles yield above chance performance). We then examined the degree to which the response retained information about texture when multiple principal components were cumulatively removed (Figure 3D). We found classification performance to be well above chance even after removing 33 principal components (95% of trial shuffles yield above chance performance). Because the outcome of these analyses may depend on the structure of the trial-to-trial variability in the response, we verified that the measured dimensionality was robust to (simulated) noise correlations (Figures S2B-D).

Finally, because PCA does not necessarily identify the most informative dimensions of response, we implemented a recently developed measure of dimensionality which is not based on explained variance (like PCA) but rather gauges the ability of the response to reliably divide up the stimulus space (cf. Rigotti et al., 2013, see Methods). Using this method, we found that the cortical response to texture can consistently classify split groups of up to 22 textures, suggesting that the texture representation in somatosensory cortex is at least 21-dimensional (Figures S2E-I). Furthermore, this measurement of 21 response dimensions is likely an underestimate: our classifier-based estimate of dimensionality is not only capped by the dimensionality of the neuronal representation, but also by the size of the stimulus set and of the recorded neuronal population (Rigotti et al., 2013). Indeed, we find that the dimensionality is still rapidly increasing as a function of neuronal group size for 141 cells (Figure S2F), so more neurons would likely yield an even higher dimensional representation in response to our texture set. In total, these classification results suggest that the dimensionality of the neural representation is driven by a large number (dozens) of components which, while often only accounting individually for a small fraction of the overall response variance, nonetheless carry significant texture information.

### Some heterogeneity in cortical responses can be attributed to differences in submodality input

Next, we examined the degree to which the cortical response inherits its structure from the periphery, where texture signals are carried by three classes of low-threshold tactile nerve fibers. To this end, we leveraged previously obtained recordings of afferent responses (from 17 SA1, 15 RA, and 7 PC fibers) to a subset of 24 textures also used in the present study (Weber et al., 2013). We then evaluated, using multiple regression, the extent to which the mean population firing rate of SA1, RA, and PC afferents evoked by these 24 common textures could account for the firing rates of individual cortical cells, using the resulting standardized regression coefficients as a gauge of the relative similarity of each tactile submodality to each cortical neuron.

First, we found that the different cortical neurons received their strongest input from different classes of tactile nerve fibers (44.7%, 37.6% and 17.7% of neurons showed maximum regression coefficients from SA1, RA, and PC afferents, respectively). Second, the responses of individual somatosensory neurons implied submodality convergence as reflected by the fact that many cortical neurons were significantly better explained by a combination of multiple afferents than they were by any single afferent (*F*-test: 28% of cells better explained by all three coefficients than any single coefficient, *p* < 0.05). Because this test has low statistical power given the small number of common stimuli between the peripheral and cortical data sets (24), we also examined the adaptation properties of cortical neurons (that is, the dynamics of their responses to trapezoidal skin indentation (Pei et al., 2009)). We found that many neurons (69%) showed both significant responses during the sustained portion of the indentation, indicative of SA1 input, as well as significant responses upon the removal of the probe, indicative of RA or PC input (Figure S3AD). Overall, 80% of neurons displayed submodality convergence by one or both of these measures. Thus, even at the single-neuron level, the texture representation in somatosensory cortex is built from signals integrated across tactile submodalities.

Next, we examined what aspects of the high-dimensional texture representation in somatosensory cortex were inherited from structure in its peripheral inputs. To this end, we recalculated our PCA on both the peripheral and cortical population responses to their shared set of 24 textures. Using canonical correlation analysis (see Methods), we found that the first three dimensions of the peripheral firing rates were significantly predictive of their cortical counterparts, but dimensions beyond these three did not yield better predictions (Figure 4A). Within this shared space, the first principal axis in cortex was highly correlated with its peripheral counterpart (*r* = 0.93). The second principal axis in cortex was also correlated with its counterpart in the periphery (r = 0.89), and this axis separated neurons with strong SA1 input (and, to a lesser extent, RA input) from those with strong PC input. Indeed, the correlation between the weight of the second principal axis in cortex and SA1, RA, and PC regression coefficient was −0.43, −0.16, 0.76, respectively. Furthermore, neurons that received strong PC input tended to produce texture responses that were correlated with each other but uncorrelated with the responses of neurons driven primarily by SA1 or RA responses (Figure 4B), reflecting the stark difference in response properties of these two sources of input. Interestingly, the most strongly PC-like cells were predominantly located in area 1 (10/12 of cells with normalized PC weight > 0.8, the other two in area 2, see Figure S1D). Thus, the second dimension of variance in the cortical response has, at one extreme, SA1-like neurons and, at the other extreme, PC-like ones. The third principal axis in cortex also showed correlation with its peripheral counterpart (r = 0.82), but its meaning is unclear. Although the first few principal axes of the texture representation in cortex are inherited from the periphery, much of the structure in the cortical representation beyond these axes cannot be explained straightforwardly from the relative strengths of SA1, RA, and PC input.

**Figure 4.**
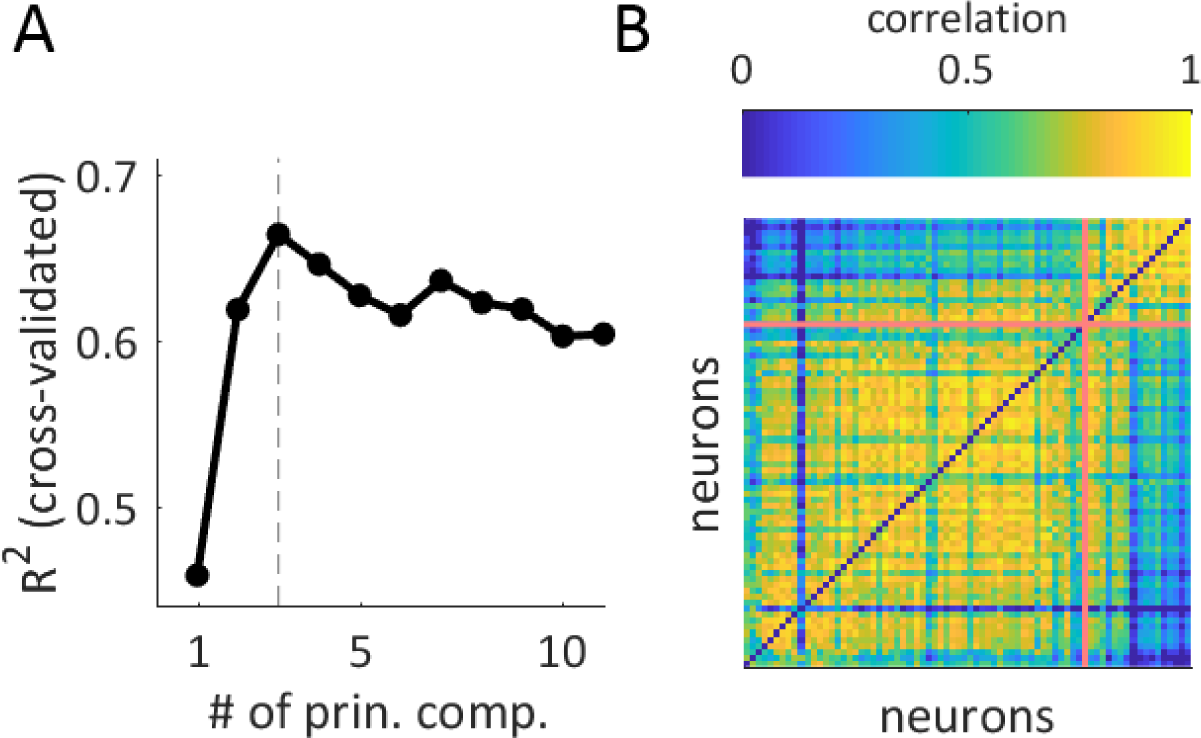
Some heterogeneity in cortical responses can be attributed to differences in submodality input. A| Strength of the prediction of cortical responses from the first N principal components of the peripheral texture response (implemented using canonical correlation analysis, see Methods). Beyond the first three principal components, performance declines due to overfitting. B| Correlation matrix of texture-elicited firing rates with each row and column corresponding to a different neuron (cells with mean texture response > 40 Hz, N=74). Cells are ordered by their PC regression weight, from least PC-like (bottom left) to most PC-like (top right). The red line divides neurons with PC regression weights greater than or less than 0.5. The most PC-like cells in somatosensory cortex tend to cluster because their texture-evoked firing rates are distinct from those of other neurons.

### Neurons in somatosensory cortex encode textural features at different spatial scales

At the periphery, texture-specific surface features are encoded through multiple mechanisms. Coarse surface features – measured in millimeters – are primarily encoded in the spatial pattern of activation across of SA1 fibers (Connor and Johnson, 1992) (and perhaps RA fibers as well (Lieber et al., 2017)). In contrast, fine surface features – typically measured in the tens or hundreds of microns – drive characteristic vibrations in the skin during texture scanning (Manfredi et al., 2014). These vibrations (and by extension, textural features) are encoded in precisely timed, texture-specific temporal patterns in RA and PC fibers (Weber et al., 2013). Next, then, we sought to examine how these peripheral codes for texture were reflected in cortical responses.

First, we tested the hypothesis that a subpopulation of somatosensory neurons act as *spatial* filters, well suited to extract information about *coarse* textural features, as has been previously proposed (DiCarlo et al., 1998; Yoshioka et al., 2001). We also wished to assess the spatial scale over which such a mechanism might operate. To this end, we first characterized the spatial receptive fields of somatosensory neurons using well-established techniques (Figures S4A-C). Using this approach, neurons have been shown to encode spatial features with excitatory subfields flanked by inhibitory ones (DiCarlo et al., 1998), analogous to simple cells in primary visual cortex (Bensmaia et al., 2008). Consistent with previous reports, the measured receptive fields exhibited well defined excitatory subfields (average 12.7 mm^2^, range 3.1-37.4 mm^2^) and inhibitory subfields (average 12.8 mm^2^, range 0-42.6 mm^2^). Inhibitory subfields tended to lag behind excitatory subfields along the scanning direction (62/67 or 93%, average 2.5 mm lag) (Figure S4D). Importantly, the spatial period of the subfield – that is, the distance between excitatory and inhibitory subfields – spanned a range from 2 to 4 mm (Figure S4E). Thus, the spatial structure of cortical receptive fields is well suited to extract information about coarse features, but not fine ones. Note that this receptive field structure is ideal for computing the spatial derivative of the neural image, which has been shown to drive perceived roughness of coarsely textured surfaces (Connor and Johnson, 1992; Lieber et al., 2017). Counterintuitively, while PC fibers have substantially larger receptive fields than do SA1 or RA fibers, this tendency was not reflected in their cortical targets. Indeed, the receptive fields of PC-like neurons were of similar size as their SA1-like or RA-like counterparts (excitatory subfield size: average 12.1 mm^2^, inhibitory subfield size: average 13.1 mm^2^, average 2.3 mm lag at 80 mm/s, see Figures S4F-G).

Next, we examined the cortical manifestation of the *temporal* code for *fine* textural features carried at the periphery by RA and PC fibers. A characteristic feature of PC (and to some extent RA) responses to texture is the elicitation of high-frequency spiking patterns (> 50 Hz) that are highly informative about texture identity, as these patterns reflect the succession of fine textural elements moving across their receptive fields (Weber et al., 2013). To explore the presence of such timing signals in the responses of somatosensory neurons, we designed two finely textured 3D patterns – gratings with spatial periods of 0.5 and 1 mm – to elicit skin vibrations at 160 and 80 Hz, respectively (given a scanning speed of 80 mm/s). We anticipated that these highly periodic components would be readily identifiable in the cortical responses, and might encode fine textural features. We found that a subpopulation of somatosensory neurons produced phase-locked responses to these and other fine textures (Figure S3E), providing a strong analog to the temporal code observed at the periphery. As expected, phase-locked responses were stronger among somatosensory neurons with PC-like responses than among their SA1-like counterparts (Figures S3E and 5B). Indeed, while the spiking patterns of both sets of neurons consistently reflected the periodic structure of coarse features, PC-like responses much more reliably reflected the periodic structure of fine features, even if these were embedded among coarse features. Neurons with PC input are thus well suited to convey information about fine textural features.

In light of these observations, we wished to assess the respective abilities of these two subpopulations of neurons – SA1-like and PC-like – to convey information about fine and coarse features. To this end, we examined the responses of these two neuronal populations to nine 3D-printed surfaces (Figure S5) in which coarse and fine features were parametrically combined (Figure 5A). We found that SA1-like neurons responded significantly more strongly to textures with coarse features than without, exhibiting only weak firing rate modulation to the presence of fine features (20.7 vs. 2.40 spikes/s for coarse vs. fine, respectively, *p*< 0.001, paired t-test). Conversely, PC-like neurons responded more strongly to textures with fine features than to those without (15.9 vs. 2.0 spikes/s, for fine vs. coarse, respectively, *p*<0.01), and their rates were nearly independent of the presence or absence of coarse features. As might be expected, these differences in sensitivity to coarse and fine textures led to corresponding differences in the ability of individual cortical neurons to discriminate pairs of textures (measured using a standard sensitivity index, *d*’). SA1-like responses were significantly better at discriminating coarse features – independent of fine features – than were their PC-like counterparts (*p*<0.05, permutation test), and PC-like neurons were significantly better at discriminating fine features – independently of the coarse features – than were SA1-like neurons (*p*<10^−4^, permutation test)(Figure 5C). In conclusion, then, different subpopulations of somatosensory neurons preferentially encode textural features at different spatial scales.

**Figure 5.**
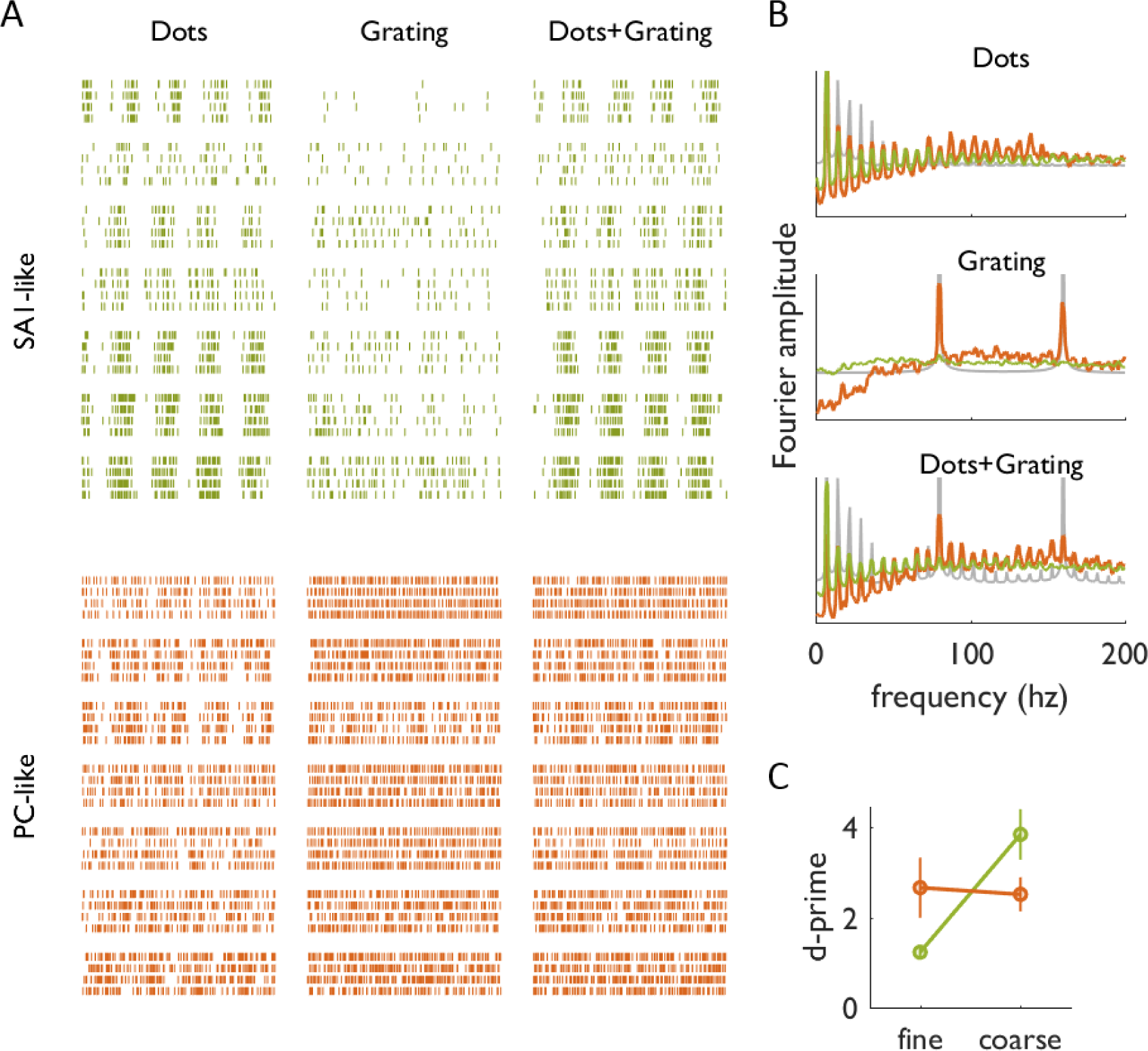
Neurons in somatosensory cortex encode textural features at different spatial scales. A| Example spiking responses are plotted from 7 SA1-like neurons (green, cells with SA1 regression coefficient > 0.5, N=53) and 7 PC-like neurons (orange, cells with PC regression coefficient > 0.5, N=23) in response to 5 repeated presentations of 3 different textures: dots spaced 7.7 mm apart, a 1 mm-period grating, and a superposition of the dots with the grating. B| Mean amplitude spectrum of the spiking responses of SA1-like (green) and PC-like cells (orange) to the same 3 textures as in A. PC-like cells exhibit high-frequency phase-locking to the temporal period of the grating (80 Hz), even when the dots are present, whereas SA1-like cells do not. C| Discriminability (*d’*) of nine 3D-printed textures based on the firing rates they evoke in SA1-like and PC-like neurons (green and orange, respectively). Error bars denote the bootstrapped standard errors of the mean across cells and texture pairs. While PC-like cells are sensitive to both coarse and fine features, SA1-like cells are sensitive only to coarse ones.

### Neuronal responses account for perceptual judgments of texture

Next, we examined how these different populations of neurons might account for the perception of texture, an important step in establishing a neural code (Blake et al., 1997; Connor and Johnson, 1992; Connor et al., 1990). To this end, we first investigated whether the responses of neurons in somatosensory cortex could account for judgments of surface roughness. Human subjects were presented with textured surfaces in an identical setup as the neurophysiological experiments, and freely rated the roughness of each surface (*n*=6 subjects, subject correlation to the mean: *r*=0.87 ± 0.079, mean ± std. dev.). We found that the firing rates of most somatosensory neurons (92%) were significantly positively correlated with roughness judgments (*r*=0.59 ± 0.27, mean + std. dev. for individual cells, 130/141 cells with significantly positive correlation at*p* <0.05, permutation test) and that the first principal component of the population response was a good predictor of roughness (Figure 6A, *r*=0.88), a consistent effect across all three cortical fields (all *r*>0.85, Figures S1E-F).

**Figure 6.**
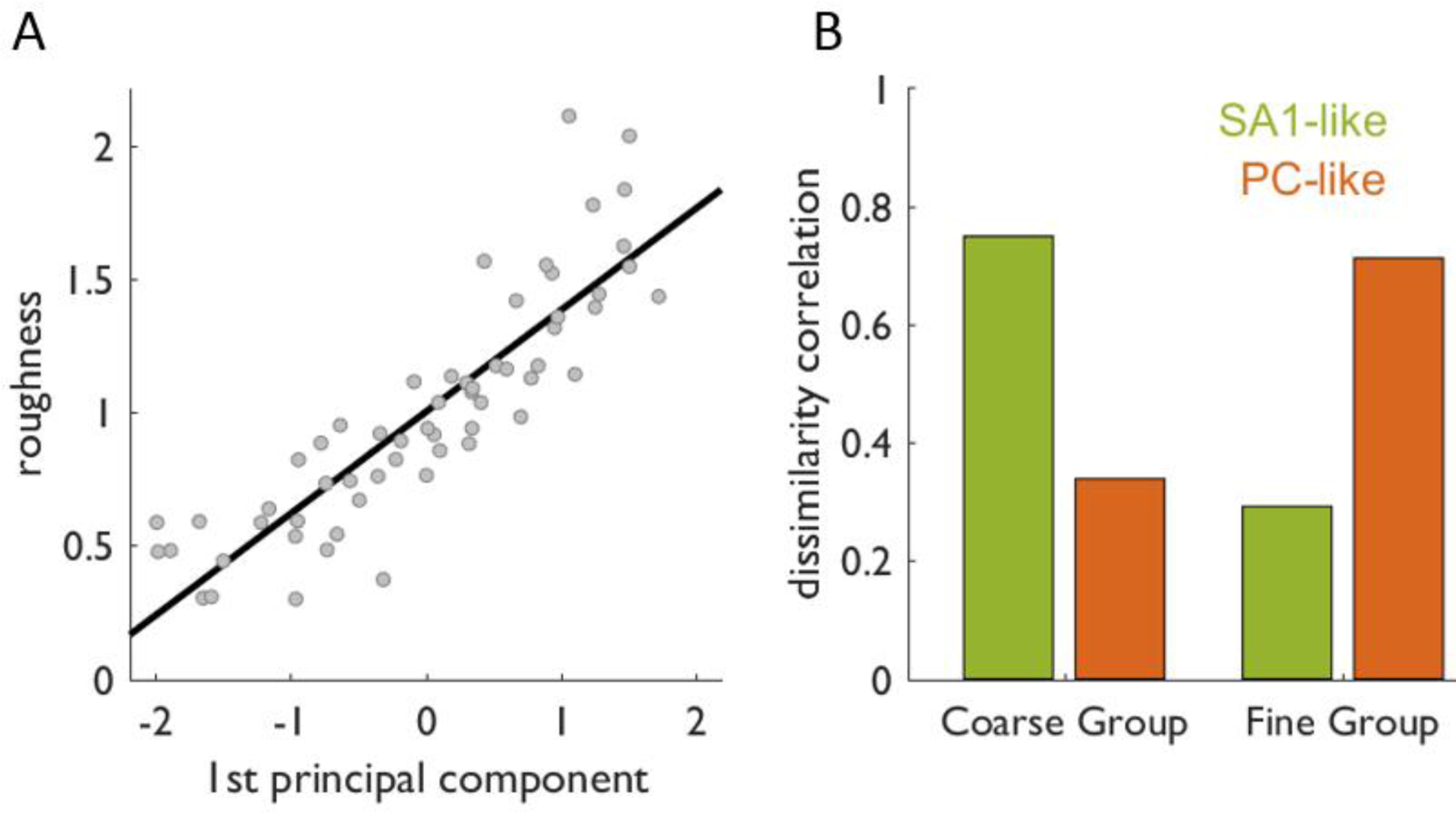
Neuronal responses account for perceptual judgments of texture. A| Perceived roughness is plotted against the first principal component of the cortical population response (*r*=0.88). B| Prediction of the perceived dissimilarity for two different groups of textures based on the firing rates of SA1-like neurons (green, SA1 coefficient > 0.5, N=53) and PC-like neurons (orange, SA1 coefficient > 0.5, N=23). SA1-like cells predict perceived differences in coarse spatial structure, whereas PC-like cells predict perceived differences in fine structure.

Although roughness is the dominant sensory dimension of texture, the perceptual space of texture also comprises other well-established sensory continua, such as hardness/softness, stickiness/slipperiness, and warmth/coolness (Hollins et al., 2000). Together, these continua combine to account for some but not all aspects of the multi-dimensional sensory experience of texture (Hollins et al., 1993, 2000; Katz, 1925). To assess the degree to which the cortical representation can account for the perceptual space, we examined the degree to which neuronal responses could account for dissimilarity judgments of texture. As with the roughness experiment, human subjects freely rated the perceived dissimilarity of pairs of textures (n=10 subjects). For this analysis, we determined the degree to which judgments of dissimilarity mirrored differences in the evoked neuronal responses (see Methods). We carried out this analysis on data obtained from two sets of texture pairs: one in which textures differed in their coarse spatial features (Coarse Group: 3 fabrics and 2 dot patterns, yielding 10 pairs, subject correlation to the mean *r*=0.88 +/- 0.10 mean ± std. dev.) and one that comprised textures mostly lacking coarse spatial features (Fine Group: 13 fabrics, 78 pairs, subject correlation to the mean *r*=0.65 ± 0.12 mean ± std. dev.) First, we examined the ability of afferent responses to predict perceived dissimilarity. We found that SA1 afferent responses best accounted for the dissimilarity of textures with different coarse spatial features (Coarse Group: correlation between firing rates and perceived dissimilarity = 0.78, 0.41, 0.17 for groups of 7 SA1, RA, and PC fibers, respectively), and PC afferent responses best accounted for the perceived dissimilarity of finely textured fabrics (Fine Group: r = 0.37, 0.39, 0.56 for groups of 7 SA1, RA, and PC fibers). When we carried out the same analysis based on cortical responses (Figure 6B), we found that SA1-like neurons best accounted for the perceived dissimilarity of Coarse Group pairs (*r* = 0.70, 0.24, 0.26 for groups of 7 SA1-like, RA-like, and PC-like neurons, respectively) and PC-like neurons best accounted for that of Fine Group pairs (*r* = 0.31, 0.26, 0.67 for groups of 7 SA1-like, RA-like, and PC-like neurons, respectively). Results from this analysis further support the hypothesis that different subpopulations of neurons encode textures at different spatial scales: SA1-like neurons are specialists for coarse textural features and PC-like neurons are specialists for fine ones.

## Discussion

### The neural mechanisms that give rise to a high-dimensional representation of texture in cortex

While texture responses in cortex are dominated by a common signal that encodes roughness, the heterogeneity of responses across individual somatosensory neurons carries considerable information about texture identity. One identifiable way in which neurons differ is in the degree to which their responses reflect SA1 afferent input vs. PC afferent input, a continuum rather than a dichotomy as evidenced by the continuous distribution of regression coefficients (Figure S1D). However, cortical responses to texture are not simply a linear combination of afferent firing rates. Indeed, while we find three shared dimensions between the peripheral and cortical representations of texture, our binary classification analysis shows that the effective number of dimensions is much higher than three (Figure 3B).

The observed increase in dimensionality from periphery to cortex is supported by an amplification of the somatosensory representation: the fingertip region of area 3b contains ∼250 cells for every corresponding peripheral afferent (Darian-Smith and Kenins, 1980; Sur et al., 1980; Turner et al., 2016). This expansion does not simply involve the establishment of redundancy in cortex, giving rise to populations of similarly tuned neurons. Rather, cortical neurons exhibit heterogeneous tuning: Each cortical neuron signals the presence of a specific spatial pattern of afferent activity within a range of spatial scales (from ∼1 to 10 mm)(DiCarlo and Johnson, 2000; DiCarlo et al., 1998) and/or a specific temporal pattern of afferent activity within a range of time scales (from ∼1 to 100 ms) (Saal et al., 2015a). While these spatial and temporal filters are often estimated using linear models, this integration of peripheral input is subject to the nonlinear motifs of neural processing, including thresholds (DiCarlo et al., 1998; Priebe and Ferster, 2012; Saal et al., 2015a), synaptic depression (Chung et al., 2002; Katz et al., 2006; Priebe and Ferster, 2012), and divisive normalization (Brouwer et al., 2015; Carandini and Heeger, 2012; Reed et al., 2010, 2011). This feature extraction and the associated non-linear transformations result in a high dimensional representation of texture, a process that is not unique to the somatosensory system. Indeed, high dimensional representations have been observed in the visual (Brincat et al., 2017; Cowley et al., 2016; Lehky et al., 2014; Stringer et al., 2018) and olfactory (Eichler et al., 2017) systems, cerebellum (Litwin-Kumar et al., 2017), hippocampus (McKenzie et al., 2014), and prefrontal cortex (Rigotti et al., 2013), to name a few.

### The perceptual space of texture is also high-dimensional

The multidimensional nature of the texture representation in somatosensory cortex reflects the complexity of the space in which surface materials and microstructures reside, and the resulting perceptual space of textures. While some aspects of this space can be captured by a small number of commonly identified sensory dimensions – roughness, hardness, stickiness, and warmth (Hollins et al., 2000) – many others cannot. That is, the roughness, hardness, stickiness, and warmth of a texture defines it only partially. Many adjectives to describe texture – fuzzy, bumpy, silky, to name just a few – evoke additional textural features not captured in low-dimensional descriptions. Further dimensions may not be simply captured by such intuitive descriptors – one can imagine fields of repeating elements arranged in different configurations that are discriminable in a way that is difficult or impossible to articulate. Thus, in a manner broadly analogous to the space of visual shape (Connor et al., 2007; Lehky et al., 2014) and visual texture (Freeman et al., 2013; Portilla and Simoncelli, 2000), the complex neural space of tactile texture is reflected in a complex perceptual space.

### Texture representations in cortex fall on a continuum of spatial scales

As described above, somatosensory neurons fall along a continuum – captured in the second principal component of their responses – that seems to be determined by their peripheral inputs. The position of a neuron along this axis relates to the spatial scale of the textures it is best suited to encode. At one end of the continuum, SA1-like neurons encode coarse textural elements; at the other end, PC-like neurons encode fine features. This differential spatial sensitivity is reflected in the ability of neurons to convey information about texture: SA1-like responses best distinguish textures with different coarse features while PC-like responses best distinguish textures with different fine features. These differences are also reflected in the ability of neuronal populations to predict perceptual judgments of texture: SA1-like neurons account for the perception of coarse features; PC-like neurons account for the perception of fine features.

In the peripheral nerve, coarse and fine textures are encoded through two mechanisms, a spatial code and a temporal one, respectively. Somatosensory neurons are well suited to extract coarse textural features measured in millimeters as evidenced by the spatial dimensions of their receptive fields (Figures S4E-F)(Bensmaia et al., 2008; DiCarlo et al., 1998). As discussed above, the idiosyncratic receptive field structure of individual neurons (Figure S4C) confers to them idiosyncratic preferences for coarse textural features and likely drives, in part, the heterogeneity of texture responses. Furthermore, the computation that such receptive fields imply – of spatial variation – has been shown to determine perceived roughness (Connor et al., 1990; Goodman and Bensmaia, 2017; Lieber et al., 2017; Weber et al., 2013). Thus, whereas *spatial variation* in afferent responses predicts roughness judgments, cortical *firing rates* predict roughness judgments (Bourgeon et al., 2016; Burton and Sinclair, 1994; Chapman et al., 2002) because they reflect the output of this differentiation computation.

At the periphery, however, the spatial mechanism cannot account for the perception of fine features due to limitations of the skin to transmit those features to the receptors (Phillips and Johnson, 1981; Sripati et al., 2006b, 2006a) and limitations set by the cutaneous innervation density (Darian-Smith and Kenins, 1980; Johansson and Vallbo, 1979). When the finger slides across a textured surface, vibrations are elicited in the skin. These vibrations are highly texture specific (Bensmaia and Hollins, 2003; Bensmaïa and Hollins, 2005; Delhaye et al., 2012; Hollins et al., 2002; Manfredi et al., 2014) and, in turn, drive precise temporal spiking patterns in RA and particularly PC fibers (Weber et al., 2013). Somatosensory neurons implement temporal variation computations (Saal et al., 2015a), which amount to extracting features in the temporal spiking patterns and converting them into rate-based signals. The nature and time scale of these temporal variation computations vary from neuron to neuron, and this heterogeneity contributes to the observed heterogeneity in texture responses. This transformation allows for the possibility, in principle, that all the relevant texture information in spike timing has been converted to a rate code in cortex.

### Is spike timing relevant to texture coding in cortex?

Though PC-like neurons in somatosensory cortex exhibit texture-specific temporal spiking patterns (Figure 5A), the role of spike patterning in texture coding remains to be elucidated. Indeed, textures in our set can be classified accurately based on the firing rates they evoke in cortical neurons, without taking into account precise spike timing. One possibility is that the texture information conveyed by spike timing in cortex is redundant with the texture information conveyed by the rates. The proposed irrelevance of spike timing to texture coding would seem to contradict its demonstrated importance for the coding of vibratory frequency (Harvey et al., 2013), given that texture-specific temporal spiking patterns are driven by skin vibrations. However, these seemingly divergent findings for vibration and texture can be reconciled by a model where temporal patterning in somatosensory cortex encodes information about the skin’s behavior, e.g. the frequency of skin vibrations (Manfredi et al., 2014), divorced of any ethological meaning. Texture perception involves the identification of spatio-temporal patterns of skin deformations produced by a meaningful stimulus, a textured surface. In this case, the relevant patterns are encoded in the rates even though the skin-related signal is still present in the temporal patterning and is perceptually available independently of the texture; for example, the periodicity of corduroy can be consciously accessed. Such a timing signal might be particularly useful for building a representation of relevant exploratory parameters (in this case scanning speed) that is tolerant to differences in surface structure (Dallmann et al., 2015). Alternatively, this timing signal may be informative for some textures, but redundant with the rate signal in our specific texture set. To establish the role of temporal signals in cortical representations of texture will require the construction of finely textured surfaces designed to draw out this temporal component.

### Conclusions

Texture representations in cortex involve the extraction of spatial and temporal features from the patterns of activation across tactile fibers. The resulting high-dimensional cortical representation of texture comprises dozens of non-redundant signals, many of which account only for a small fraction of the overall neuronal variance but are nonetheless informative about texture identity. A prominent axis in the neuronal response forms a continuum of spatial scales, with coarse feature specialists at one extreme and fine feature specialists at the other, each population receiving dominant input from a different class of tactile fibers. This structure in the neuronal representation is reflected in perceptual judgments of textures. While the principal neural axis predicts perceived roughness, the code for texture identity seems to be distributed along the neural continuum of spatial scales. That is, coarse feature specialists predict coarse texture perception and fine feature specialists predict fine texture perception.

## Acknowledgements

We would like to thank Alison Weber and Ju-Wen Cheng for collecting the peripheral nerve data, Frank Dammann, Michael Harvey, Oksana Lasowsky and Erik Schluter for assistance in setting up the cortical experiments, and Benoit Delhaye, Katie Long, Hannes Saal, and Jeffrey Yau for comments on a previous version of the manuscript. This work was supported by NINDS RO1 NS101325.

## Author Contributions

J.D.L. and S.J.B. designed the research; J.D.L. performed the research and analyzed the data; J.D.L. and S.J.B. wrote the paper.

## Methods

### Experimental methods

#### Behavioral training

Before the beginning of recording sessions, all animals were trained to sit in a primate chair with their heads fixed and arms restrained as they were habituated to the experimental apparatus. During the task, the arm was stabilized in a supinated position with a custom-built cast (Polycaprolactone, lined on the interior with foam padding for comfort). The animal was trained to keep its hand still for the duration of the recording protocols, and a protocol was restarted from the beginning if the finger moved. Stability was further maintained by loosely taping the non-stimulated fingers down, and applying a small amount of glue to the fingernail of the stimulated finger to keep it in stable contact with the hand holder.

To maintain alertness during recording, the animals performed a simple visual brightness discrimination task (Harvey et al., 2013). Briefly, the animals fixated on a small square presented in the center of a monitor located in front of the tactile stimulator. After approximately 1-2s of fixation, two circles of different luminance appeared to the left and right of the fixation point. The animal was given a liquid reward (juice or water, depending on the animal’s preference) for making a saccade to the brighter target. The task was kept challenging by adjusting the relative luminance of the targets and the fixation time. Eye movements were tracked using a camera-based eye tracker (ViewPoint PC-60, Arrington Research, Scottsdale, AZ) and visual stimuli were presented using in-house software based on the OpenGL library.

#### Surgery

Procedures were approved by the University of Chicago Institutional Animal Care and Use Committee. First, a custom-built head-post was secured to the skull and allowed to osseointegrate for 1.5 months before the head was first immobilized. Once the animals were sufficiently habituated to the test apparatus and visual task, a recording chamber (22 mm internal diameter) was attached to the skull using bone cement such that it circumscribed the hand representation in somatosensory cortex, and a craniotomy was made over the internal diameter of the chamber. All surgical procedures were performed under sterile conditions; anesthesia was induced with ketamine and dexmedatomadine, and maintained with a surgical plane of isofluorane and occasional redosing of dexmedatomadine (Theriault et al., 2008). Post-surgery, anesthesia was reversed with atipamezole.

#### Neurophysiological procedures

Extracellular recordings were made in the postcentral gyri of three hemispheres of three macaque monkeys (male, 6-8 yrs old, 8-11 kg) using previously described techniques (Harvey et al., 2013). On each recording day, a multielectrode microdrive (NAN Instruments) was loaded with three tungsten electrodes insulated with epoxylite (FHC Inc., Bowdoin, ME) and electrodes were lowered normal to the cortical surface, through a custom-designed 3D-printed guide tube system that arranged the electrodes in a line 650 microns apart tip-to-tip. The electrodes were then driven into the cortex until they encountered neurons from areas 3b, 1, and 2 of somatosensory cortex with RFs on the distal fingerpad.

The transition from area 1 to area 3b exhibits a characteristic progression of RF locations. As one descends from the cortical surface through area 1 into area 3b near the central sulcus, the RFs progress from the medial and proximal finger pads to the palmar whorls. As one enters area 3b, RFs proceed back up the finger, transitioning from proximal, to medial, and ultimately to distal pads. Because responses from the distal pad were never encountered in the more superficial regions of 3b (where the palmar whorls or proximal pad typically were most responsive), there was never any uncertainty about the anatomical area from which area 3b recordings originated. The representation of the digits in area 2 lies just caudal to, and mirrors that of area 1. Thus, as one proceeds caudally, one first encounters the proximal, then medial, then distal pads. As one enters area 2, RFs remain on the distal pads and then proceed down the finger as one further proceeds caudally. The most salient feature identifying area 2 is the presence of neurons with proprioceptive response properties; that is, neurons that respond preferentially to movements of the joints. Because the distal finger pad representations in areas 1 and 2 are adjacent, we used the presence of proprioceptive responses to inform the areal classification.

We recorded from neurons whose RFs were located on the distal pads of digits 2–5. On roughly every second day of recording, the electrode array was shifted 200 mm along the postcentral gyrus until the entire representation of digits 2–5 had been covered. At the end of the recording day, the electrodes were withdrawn and the chamber was filled with sterile saline and sealed. Recordings were obtained from neurons in areas 3b, 1, and 2 that met the following criteria: (1) action potentials were well isolated from the background noise, (2) the RF included at least one of the distal finger pads on digits 2-5, (3) the finger could be positioned such that the textured surface impinged on the center of the RF, and (4) the neuron was clearly driven by light cutaneous touch. Isolations had to be maintained for at least 30 minutes to complete 5 repetitions of the basic texture protocol. When held for longer, additional protocols were run (see below).

#### Stimulus presentation

Textured surfaces were presented to the fingertips of awake macaque monkeys using a custom-built rotating drum stimulator like those used in previous studies (Johnson and Phillips, 1988; Weber et al., 2013), but larger and more precise. The drum was attached to a rotation motor (SmartMotor SM23165D, Animatics, Columbus, OH) via a 1:100 gearbox (Animatics), which provided precise control of rotational position (± 200 µm) and velocity (± 1.1 mm/s). The motor was attached to a vertical stage (IMS100V, Newport, Irvine, CA), which could control the depth of indentation into the skin with a precision of 2 microns. The vertical stage was attached to another horizontal stage (IMS400CCHA, Newport) allowing smooth displacement over 40 cm at a precision of 4 microns. Thus, we achieved precise horizontal, vertical, and rotational positioning of textures, allowing 60 different slots (12 rows, 5 textures per row) in which texture strips (2.5 cm wide by 16 cm long) could be mounted to the drum (25.5 cm in diameter and 30 cm in length). The inter-stimulus interval was at least 3 s between stimulus presentations, to allow the drum to reposition and to prevent neural adaptation.

One of the available slots was dedicated to a small load cell placed on the surface of the drum (LSB200, Futek, Irvine, CA; 2 lb, 1-axis, parallel to indentation), whose function was to ensure a consistent level of pressure was exerted on the finger across recording sessions. Each day, after the animal’s hand and finger were stabilized in place, the drum was rotated and translated such that the load cell was pressed lightly (to a force of 15 ± 3 g) into the animal’s fingertip. This position was used as a reference point for all protocols, to ensure that stimulation was consistent across days and across fingers.

#### Stimuli

Texture samples were mounted on individual strips of magnetic tape (5 × 16 cm), which were then attached to a complementary sheet of magnetic tape fixed to the surface of the drum. This allowed for simple removal and replacement when textures were damaged or worn through use. In total, 59 different textures were mounted on the drum, including a wide array of natural textures such as papers, fabrics, furs, and upholsteries with coarse periodic structure, as well as tetragonal arrays of embossed dots (Connor et al., 1990), and 3D-printed gratings and dots. Twenty four of these textures were also used in a previous experiment on peripheral afferents (Weber et al., 2013) (see Table S1).

Textures were presented at a speed of 80 mm/s and at a force of 15 g. To find the displacement equivalent to this desired force, a set of calibration readings were taken offline using a 2^nd^ load cell mounted at the location of the hand. First, a standard reference point was found by indenting the drum-mounted load cell into the 2^nd^ load cell, to a force of 15 g. Then, individual textures were repeatedly indented into the 2^nd^ load cell to find the displacement (relative to the calibration point) necessary to achieve the calibration force (15 g). During recording, textures were scanned over the finger according to these standard displacements relative to the reference point, measured daily.

#### Peripheral recordings

We have previously reported the responses of 35 afferent fibers (15 SA1, 13 RA, and 7 PC, characterized using standard criteria based on their response properties (Muniak et al., 2007)) to 55 different texture stimuli (see ref. (Weber et al., 2013) for details). Briefly, we collected extracellular single-unit recordings from the median and ulnar nerves of six anesthetized (isoflurane) Rhesus macaques as texture stimuli were presented to the distal digits of the hand at a speed of 80 mm/s. Each texture was presented to each fiber at least twice.

#### Adaptation protocol

One way to assess the submodality composition of a neuron is to measure its response to a skin indentation. Indeed, SA1 fibers are the only ones to respond during the sustained portion of the indentation whereas RA and PC fibers are the only ones to respond during the offset of the indentation. To the extent that a cortical neuron exhibits both of these properties, we can infer that it receives convergent input from multiple classes of tactile fibers. With this in mind, we measured the response of cortical neurons to a probe indented into the skin. Specifically, we indented the drum-mounted load cell (a cylinder of diameter 7 mm, see above) vertically into the skin to the 15 g reference point, and held the indentation for 500 ms, before removing it from the skin (5 mm/s indentation/removal speed). This indentation was repeated 60 times, with an inter-trial interval of 500 ms. We completed this protocol for 94 neurons (28, 57, and 9 from areas 3b, 1, and 2, respectively).

#### Spatial receptive fields

To systematically measure the spatial receptive field of individual cortical neurons, we designed a 3D-printed random dot array following a previously described approach (DiCarlo et al., 1998) (truncated cones: 0.5 mm dot height, 1 mm base diameter, 0.5 mm top diameter, 10 dots/cm^2^, dots uniformly distributed, 5 cm × 16 cm). The random dot pattern was scanned over the skin 100 times but, after each scan, the drum was stepped 400 microns along its axis of rotation. A full uninterrupted protocol was completed (∼10 minutes), for 72 neurons (27, 37, and 8 from areas 3b, 1, and 2, respectively).

#### Magnitude estimation

All procedures were approved by the Institutional Review Board of the University of Chicago and all subjects provided informed consent. Subjects sat with the right arm supinated and resting on a support under the drum. Stimuli were presented to the right index fingerpad of each subject.

*Roughness scaling*(6 subjects, 5m, 1f, ages 18-24): On each trial, the subject was presented with one of 59 textures (80 mm/s, 25 ± 10 g) and produced a rating proportional to its perceived roughness, where a rating of zero denoted a perfectly smooth surface. If texture B was perceived to be twice as rough as texture A, texture B was ascribed a number that was twice as large as texture A. Subjects were encouraged to use fractions and decimals if necessary. Each texture was presented once in each of 6 experimental blocks; ratings were normalized by the mean of each block and averaged, first within then across subjects. Ratings of roughness were consistent across subjects (subject correlation to the mean: *r*=0.87 ± 0.079, mean ± std. dev.).

*Dissimilarity scaling* (10 subjects, 10f, ages 19-24): In these experiments, two subsets of textures were used. The first comprised 5 textures (Coarse Group: silk, microsuede, upholstery, 4 mm embossed dots, 5 mm embossed dots), and the second 13 textures (Fine Group: suede, chiffon, nylon (200 denier), denim, hucktowel, silk, microsuede, wool, satin, metallic silk, upholstery, thin corduroy, thick corduroy). On each trial, the subject was presented with a pair of textures (for 1 s each, separated by a 1 s inter-stimulus interval, 10/78 unique comparisons for each group, respectively, 80 mm/s, 25 ± 10 g) and produced a rating proportional to the perceived dissimilarity of the pair, where 0 denotes (perceived) identicalness. Each pair of textures was presented 3 times in pseudorandom order. Texture dissimilarity ratings were correlated across subjects (subject correlation to the mean, Coarse Group: *r*=0.88 +/- 0.10, Fine Group: 0.65 ± 0.12, mean ± std. dev.).

### Analysis

#### Basic analyses of firing rate

Because the stimulus epoch over which a texture was moved into the skin evoked a large phasic response that lasted about 200 ms, we excluded this response from our analysis. For each trial, the baseline firing rate was measured in the 500-ms period before drum’s initial contact with the skin.

To test whether textures responses were significantly above baseline firing rates, we first created a distribution of baseline responses for each cell. We then measured how often each cell’s texture firing rate response (averaged across 5 repetitions) was greater than the average of 5 random draws from the baseline distribution. We set significance at p<0.05 for all textures, Bonferroni corrected for 8319 comparisons (141×59).

For data from the indented probe protocol, we measured the trial-by-trial firing rate over stimulus epochs: 1) background activity, measured between 1 s and 500 ms before the probe was indented into the skin, 2) sustained response activity, measured during the 500 ms hold period, and 3) offset response activity, measured between 100 and 300 ms after the probe started lifting off the skin. We report that a neuron has a significant sustained/offset response if its texture elicited firing rate was significantly different from baseline (two-sided t-test for sustained/offset-baseline, significant if *p*<0.05). To quantify the relative magnitude of the sustained and offset responses, we calculated the fraction of the combined sustained and offset responses that was carried by the offset response:

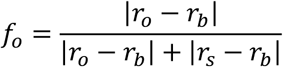

where *r_o_* is the firing rate at the offset, *r_s_* is the firing rate during the sustained phase, and *r_b_* is the baseline firing rate.

#### Principal components analysis

We applied a principal components analysis (PCA) to population responses to determine their major axes of variation. Specifically, we treated each cortical neuron as a signal, and each texture response as an observation in 141-dimensional space.

To compare the main axes of variation in the responses of tactile nerve fibers and cortical neurons, we identified shared axes of variation using a cross-validated canonical correlation analysis. First, we split the 24 textures that were used in both the peripheral and cortical experiments into a “training” set of 23 textures, leaving out 1 “test” texture. Second, we recalculated the PCA to determine new axes of variation for the peripheral (N = 39) and cortical (N = 141) responses. Third, we used canonical correlation analysis (using the *canoncorr()* function in Matlab) to calculate the optimal mapping of component scores from the peripheral to the cortical representation. Fourth, we used this mapping on the responses to the “test” texture: specifically, we applied the mapping to the afferent population response and obtained from it a prediction of the cortical population response. This procedure was repeated 24 times, each time leaving one texture out of the training set, and was repeated with different numbers of principal axes (between 1 and 22 principal components). Finally, for each set of principal axes, we computed the coefficient of determination for this cross-validated procedure, using a formula analogous to the traditional *R^2^* value for linear regression:

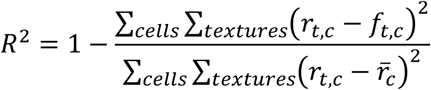

where *r_t, c_* is the true firing rate of a cell *c* in response to the test texture *t*, *f_t, c_* is the predicted firing rate of cell *c* in response to that texture, and 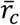 is the mean firing rate of cell *c* averaged across the 24 textures.

#### Texture classification

To assess the degree to which neuronal responses carry information about texture identity, we assessed the degree to which we could classify textures based on the neuronal responses they evoke. To this end, we implemented a nearest neighbor classifier. First, for each neuron, we averaged 4 of the 5 responses evoked by each texture, leaving 1 repetition out, yielding 59 vectors of mean responses and 59 vectors of single-trial responses. Next, we computed the distance between each single-trial response and each mean response. For each single trial response, the mean response yielding the lowest distance was selected. If the selected mean responses and the single trial responses corresponded to the same texture, classification was correct. Performance was averaged across all textures, and then again across 100 shuffles of which repetitions were left out. This procedure was repeated for neuron groups of different sizes.

We wished to examine how well responses from the cortical population could support texture classification using subsets of the full response space, as defined by the axes of variation identified via PCA. To implement our classification analysis in these subspaces, we first recomputed the PCA using the mean firing rates computed from only 4 repetitions (the training set), as discussed above. Next, we projected the full response space (for the training and test sets) onto the relevant subspace (either single dimensions, as in Figures 3C and S2C, or lower-dimensional subspaces, as in Figures 3D and S2D). Finally, we performed the classification, as described above, in this lower-dimensional subspace.

For one analysis, we performed a pairwise texture classification, rather than the “one in 59” classification described above (Figures 3C and S2C). Classification in this case was performed exactly as described above, but only for comparing the responses to two textures at a time, rather than comparing any one texture to the remaining full set of textures. Performance was averaged over all possible pairs of textures, and then again over all “test” repetitions.

#### Binary classification and dimensionality

The dimensionality of the cortical population response estimated from PCA comprises components that reliably carry texture information and components that do not. To estimate the components of the dimensionality that contribute to the texture representation, we adapted a method from (Rigotti et al., 2013) that assesses the number of informative dimensions by quantifying the ability of the population representation to perform arbitrary linear binary classifications of the state space (in this case textures). The idea is, to the extent that the texture set can be arbitrarily split into *D* categories, the neural space comprises *D* dimensions relevant to texture coding. First, we randomly select N neurons and T textures. We choose one (of the 2^T^ possible ways) of splitting the T textures into two groups, and test whether a binary classifier could successfully discriminate between the two groups using the N-dimensional population response. Specifically, we train a support vector machine with a linear kernel (with the *fitcsvm()* function in Matlab) on four randomly selected responses to each texture (out of five, without replacement), each yielding an N-dimensional vector of firing rates. We then use the remaining left-out response from each cell to build a single N-dimensional test vector for each texture, which is used to test the performance of the binary classifier. Performance is averaged over 50 repetitions of the trial-shuffling procedure, to get a mean performance for this set of cells and texture groups. If this mean performance is greater than 75%, we count this binary classification as “implementable” by the population response. The functions in Figure 3B represent the proportion of “implementable” conditions, measured over 500 random selections of cells and texture groupings. Data are fit with sigmoids:

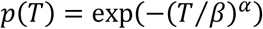

and the intersection of this curve *p(T)* = 0.95 is taken as the critical value T* for each population size. A population that can successfully classify T* textures has an effective dimensionality of T*-1. Thus, in Figure 3C, we report the dimensionality for a population size N as *D = max(T*−1, 0)*.

This method of measuring dimensionality provides a lower bound on the true dimensionality of the neural representation (Rigotti et al., 2013). To verify this, we generated representations of texture with known dimensionality, and then measured the dimensionality of these new texture responses using the binary classification metric (see Figures S2E-I). Specifically, to create an N-dimensional space of texture responses, we constructed new responses by deriving each neuron’s mean firing rate from the first N principal components. To obtain single trial responses, we subtracted the raw mean firing rate for each texture from each single-trial response and added the corresponding mean reconstructed from the lower-dimensional representation. Thus, the trial-to-trial variability in the responses of individual neurons to individual textures was preserved, but the dimensionality of the population response was reduced to N.

#### Simulating noise correlations

We sought to test the impact on texture classification and neural dimensionality of noise correlations – absent from neuronal responses because they were measured non-simultaneously – by simulating the effects of noise correlations using a resampling technique. First, for each texture, we generated a set of 5 population response vectors (across all 141 cells) using data from its 5 repeated presentations. These repetitions were ordered from highest to lowest firing rate, such that the first population vector contained the highest firing rate response for each individual cell, the second vector contained the second highest rate response, etc. Repeating this procedure across all textures created a set of population vectors (5×59) with a very high level of mean noise correlation (*r_sc_*=0.74). To bring this value to physiologically relevant levels (*r_sc_*=0.1-0.2 (Cohen and Kohn, 2011)), we repeatedly looped over each cell/texture combination (10 times) and randomly swapped two of the repetitions between population vectors with a set probability (*p* = 0.15, 0.2, 0.26 and 0.8). This resulted in sets of population vectors with physiologically relevant levels of mean noise correlation (*r_sc_*=0.22, 0.14, 0.08 and 0.001, respectively).

To perform our texture classification with these resampled neuronal responses, we repeated the classification procedure described above with only one difference: performance was only calculated using each of the 5 population vectors with targeted correlations as the “single trial response,” rather than using the randomly selected responses on each shuffle. When applicable, responses were still averaged over 100 randomly selected groups of cells at each group size. We also recalculated our dimensionality analyses using these resampled data sets. Rather than using the random shuffling procedure, we iterated over each of the 5 simulated population vectors as the “left out response.” Each point in Figure S2G represents the mean number of “implementable” texture sets, averaged over 300 random texture groupings.

#### Estimating submodality input

We wished to assess the relative contributions of the three functionally-defined populations of tactile fibers to the response of each neuron in somatosensory cortex, having previously shown that a majority of cortical neurons receive convergent input from multiple modalities, even in area 3b (Pei et al., 2009; Saal et al., 2015b) (recognizing that afferent signals pass through at least two intermediate synapses, one in the cuneate nucleus and one in the thalamus). While afferent input is likely not integrated linearly, we estimated the relative strength of that input using a linear model. Specifically, we used a multiple regression to predict the standardized (z-scored) mean texture responses of each cortical neuron to a set of 24 textures (see above) from the standardized (z-scored) mean responses of SA1, RA, and PC afferents to those same textures. We used these normalized regression weights as measures of the relative strength of SA1, RA, and PC afferent input into each neuron.

#### Spatial receptive fields

To characterize the spatial receptive field of each cortical neuron, we adopted an approach described in detail in ref. (DiCarlo et al., 1998). In brief, we first binned spiking responses to the receptive field protocol (see above) in 100 micron (1.25 ms) bins. The 100 runs – each corresponding to the response during one scan of the random dot pattern, each scan radially displaced from the previous one by 400 microns – were combined into a neural image (spatial even plot, cf. ref. (Johnson and Phillips, 1988)), which was then cross-correlated with a reconstruction of the stimulus (binned into 100×100 micron bins) to find an optimal alignment. Then, we used a spike-triggered average (STA) of the stimulus to find the mean stimulus values that were spatially aligned to any given spike.

To calculate receptive field properties, we first smoothed the STA with a 0.3 mm std. dev. Gaussian filter. Next, all bins with an absolute amplitude less than 20% of the RF’s peak amplitude were set to zero. Finally, we required 1) that every non-zero bin have at least two of the four adjacent bins be nonzero and 2) that isolated regions of the RF have areas of at least 0.7 mm^2^. Bins that did not meet these criteria were set to zero. To calculate excitatory & inhibitory area, we summed the area occupied by bins in the RF with positive/negative values, respectively. To calculate the scanning distance between excitatory and inhibitory subfields, we first found the center-of-mass for the excitatory and inhibitory bins in each RF, and then calculated the distance between them along the scanning direction.

#### Frequency analysis

Spiking responses in the nerve have been shown to phase-lock at high frequencies to texture-elicited skin vibrations (Weber et al., 2013), and spiking responses in somatosensory cortex have been shown to phase-lock to high-frequency skin vibrations imposed by a vibrating probe (Harvey et al., 2013). To reveal any phase-locking in the cortical responses to texture, we first binned spike trains into 0.3 ms bins. Next, we computed the fast Fourier transform of that binned spike train and, from it, the amplitude spectrum. Finally, we computed the mean amplitude spectrum across repeated presentations for each texture and neuron.

#### Discriminability of 3-D printed textures

We wished to assess whether neurons in somatosensory cortex are more sensitive to differences in the coarse structure or fine structure of a textured surface. To this end, we measured neuronal responses to nine 3D-printed textures that parametrically combined three coarse patterns (blank, 7.7 mm spaced dots, 5 mm-period grating) and three fine patterns (blank, 1 mm-period grating, and 500 µm-period grating). We then measured the discriminability of pairs of textures from their respective distributions of responses. Specifically, we sought to determine whether two textures with identical fine structure, but different coarse structure could be discriminated from the responses of an individual neuron. For each fine pattern, we had three different coarse patterns (and vice versa for each coarse pattern), and thus 3 different pairwise comparisons. To quantify the discriminability of each pair from each neuron’s responses, we measured a sensitivity index (*d’*):

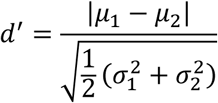

where *μ* is the mean firing rate response to a texture and *σ^2^* is the variance in firing rate across repeated presentations of that texture. These values were averaged across conditions and subpopulations of cells (Figure 5C). To calculate the statistical significance of these differences across neural subpopulations, we repeated this averaging on permutations the same data set with responses shuffled (50000 permutations). We calculated significance as the proportion of times our measured difference was greater than that computed from shuffled data.

#### Dissimilarity correlation

We sought to determine whether we could account for psychophysical judgments of texture dissimilarity obtained from human observers to neuronal responses in somatosensory cortex. To this end, we computed the Euclidean distance between the two population vectors for each of the two textures, each element of which is the mean firing rate of each neuron in the set. This distance was then compared with the psychophysical judgment of texture dissimilarity.

## References

Bensmaia, S.J., and Hollins, M. (2003). The vibrations of texture. Somatosens. Mot. Res. 20, 33–43.

Bensmaia, S.J., Denchev, P. V., Dammann, J.F., Craig, J.C., and Hsiao, S.S. (2008). The representation of stimulus orientation in the early stages of somatosensory processing. J. Neurosci. 28, 776–786.

Bensmaïa, S.J., and Hollins, M. (2005). Pacinian representations of fine surface texture. Percept. Psychophys. 67, 842–854.

Blake, D.T., Johnson, K.O., and Hsiao, S.S. (1997). Monkey cutaneous SAI and RA responses to raised and depressed scanned patterns: effects of width, height, orientation, and a raised surround. J. Neurophysiol. 78, 2503–2517.

Bourgeon, S., Dépeault, A., Meftah, E.-M., and Chapman, C.E. (2016). Tactile texture signals in primate primary somatosensory cortex and their relation to subjective roughness intensity. J. Neurophysiol. 115, 1767–1785.

Brincat, S., Siegel, M., von Nicolai, C., and Miller, E.K. (2017). Gradual progression from sensory to task-related processing in cerebral cortex. BioRxiv 195602.

Brouwer, G.J., Arnedo, V., Offen, S., Heeger, D.J., and Grant, A.C. (2015). Normalization in human somatosensory cortex. J. Neurophysiol. 114, 2588–2599.

Burton, H., and Sinclair, R.J. (1994). Representation of tactile roughness in thalamus and somatosensory cortex. Can. J. Physiol. Pharmacol. 72, 546–557.

Carandini, M., and Heeger, D.J. (2012). Normalization as a canonical neural computation. Nat. Rev. Neurosci. 13, 51–62.

Chapman, C.E., Tremblay, F., Jiang, W., Belingard, L., and Meftah, E.M. (2002). Central neural mechanisms contributing to the perception of tactile roughness. In Behavioural Brain Research, pp. 225–233.

Chung, S., Li, X., and Nelson, S.B. (2002). Short-term depression at thalamocortical synapses contributes to rapid adaptation of cortical sensory responses in vivo. Neuron 34, 437–446.

Cohen, M.R., and Kohn, A. (2011). Measuring and interpreting neuronal correlations. Nat. Neurosci. 14, 811–819.

Connor, C.E., and Johnson, K.O. (1992). Neural coding of tactile texture: comparison of spatial and temporal mechanisms for roughness perception. J. Neurosci. 12, 3414–3426.

Connor, C.E., Hsiao, S.S., Phillips, J.R., and Johnson, K.O. (1990). Tactile roughness: neural codes that account for psychophysical magnitude estimates. J. Neurosci. 10, 3823–3836.

Connor, C.E., Brincat, S.L., and Pasupathy, A. (2007). Transformation of shape information in the ventral pathway. Curr. Opin. Neurobiol. 17, 140–147.

Cowley, B.R., Smith, M.A., Kohn, A., and Yu, B.M. (2016). Stimulus-Driven Population Activity Patterns in Macaque Primary Visual Cortex. PLoS Comput. Biol. 12.

Dallmann, C.J., Ernst, M.O., and Moscatelli, A. (2015). The role of vibration in tactile speed perception. J. Neurophysiol. 114, 3131–3139.

Darian-Smith, I., and Kenins, P. (1980). Innervation density of mechanoreceptive fibres supplying glabrous skin of the monkey’s index finger. J. Physiol. 309, 147–155.

Darian-Smith, I., Sugitani, M., Heywood, J., Karita, K., and Goodwin, A.W. (1982). Touching textured surfaces: cells in somatosensory cortex respond both to finger movement and to surface features. Science 218, 906–909.

Delhaye, B., Hayward, V., Lefèvre, P., and Thonnard, J.-L. (2012). Texture-induced vibrations in the forearm during tactile exploration. Front. Behav. Neurosci. 6, 37.

DiCarlo, J.J., and Johnson, K.O. (2000). Spatial and temporal structure of receptive fields in primate somatosensory area 3b: effects of stimulus scanning direction and orientation. J. Neurosci. 20, 495–510.

DiCarlo, J.J., Johnson, K.O., and Hsiao, S.S. (1998). Structure of receptive fields in area 3b of primary somatosensory cortex in the alert monkey. J. Neurosci. 18, 2626–2645.

Eichler, K., Li, F., Litwin-Kumar, A., Park, Y., Andrade, I., Schneider-Mizell, C.M., Saumweber, T., Huser, A., Eschbach, C., Gerber, B., et al. (2017). The complete connectome of a learning and memory centre in an insect brain. Nature 548, 175–182.

Freeman, J., Ziemba, C.M., Heeger, D.J., Simoncelli, E.P., and Movshon, J.A. (2013). A functional and perceptual signature of the second visual area in primates. Nat. Neurosci. 1–12.

Goodman, J.M., and Bensmaia, S.J. (2017). A variation code accounts for perceived roughness. Sci. Rep.

Harvey, M.A., Saal, H.P., Dammann, J.F., and Bensmaia, S.J. (2013). Multiplexing Stimulus Information through Rate and Temporal Codes in Primate Somatosensory Cortex. PLoS Biol. 11.

Hollins, M., and Risner, S.R. (2000). Evidence for the duplex theory of tactile texture perception. Percept. Psychophys. 62, 695–705.

Hollins, M., Faldowski, R., Rao, S., and Young, F. (1993). Perceptual dimensions of tactile surface texture: A multidimensional scaling analysis. Percept. Psychophys. 54, 697–705.

Hollins, M., Bensmaia, S., Karlof, K., and Young, F. (2000). Individual differences in perceptual space for tactile textures: evidence from multidimensional scaling. Percept. Psychophys. 62, 1534–1544.

Hollins, M., Bensmaïa, S.J., and Washburn, S. (2001). Vibrotactile adaptation impairs discrimination of fine, but not coarse, textures. Somatosens. Mot. Res. 18, 253–262.

Hollins, M., Bensmaia, S.J., and Roy, E.A. (2002). Vibrotaction and texture perception. Behav. Brain Res. 135, 51–56.

Johansson, R., and Vallbo, Å. (1979). Tactile sensibility in the human hand: relative and absolute densities of four types of mechanoreceptive units in glabrous skin. J. Physiol. 283–300.

Johnson, K., and Lamb, G. (1981). Neural mechanisms of spatial tactile discrimination: neural patterns evoked by braille-like dot patterns in the monkey. J. Physiol. 117–144.

Johnson, K.O., and Phillips, J.R. (1988). A rotating drum stimulator for scanning embossed patterns and textures across the skin. J. Neurosci. Methods 22, 221–231.

Katz, D. (1925). The World of Touch.

Katz, Y., Heiss, J.E., and Lampl, I. (2006). Cross-whisker adaptation of neurons in the rat barrel cortex. J. Neurosci. 26, 13363–13372.

Lehky, S.R., Kiani, R., Esteky, H., and Tanaka, K. (2014). Dimensionality of Object Representations in Monkey Inferotemporal Cortex. Neural Comput. 26, 2135–2162.

Lieber, J.D., Xia, X., Weber, A.I., and Bensmaia, S.J. (2017). The Neural Code for Tactile Roughness in the Somatosensory Nerves. J. Neurophysiol. jn.00374.2017.

Litwin-Kumar, A., Harris, K.D., Axel, R., Sompolinsky, H., and Abbott, L.F. (2017). Optimal Degrees of Synaptic Connectivity. Neuron 93, 1153–1164.e7.

Manfredi, L.R., Saal, H.P., Brown, K.J., Zielinski, M.C., Dammann, J.F., Polashock, V.S., and Bensmaia, S.J. (2014). Natural scenes in tactile texture. J. Neurophysiol. 111, 1792–1802.

McKenzie, S., Frank, A.J., Kinsky, N.R., Porter, B., Rivière, P.D., and Eichenbaum, H. (2014). Hippocampal representation of related and opposing memories develop within distinct, hierarchically organized neural schemas. Neuron 83, 202–215.

Muniak, M.A., Ray, S., Hsiao, S.S., Dammann, J.F., and Bensmaia, S.J. (2007). The neural coding of stimulus intensity: linking the population response of mechanoreceptive afferents with psychophysical behavior. J. Neurosci. 27, 11687–11699.

Pei, Y.-C., Denchev, P. V., Hsiao, S.S., Craig, J.C., and Bensmaia, S.J. (2009). Convergence of submodality-specific input onto neurons in primary somatosensory cortex. J. Neurophysiol. 102, 1843–1853.

Phillips, J., and Johnson, K. (1981). Tactile spatial resolution. II. Neural representation of bars, edges, and gratings in monkey primary afferents. J. Neurophysiol. 1192–1203.

Portilla, J., and Simoncelli, E.P. (2000). A Parametric Texture Model Based on Joint Statistics of Complex Wavelet Coefficients. Int. J. Comput. Vis. 40, 49–71.

Priebe, N.J., and Ferster, D. (2012). Mechanisms of Neuronal Computation in Mammalian Visual Cortex. Neuron 75, 194–208.

Reed, J.L., Qi, H.-X., Zhou, Z., Bernard, M.R., Burish, M.J., Bonds, A.B., and Kaas, J.H. (2010). Response properties of neurons in primary somatosensory cortex of owl monkeys reflect widespread spatiotemporal integration. J. Neurophysiol. 103, 2139–2157.

Reed, J.L., Qi, H.-X., and Kaas, J.H. (2011). Spatiotemporal Properties of Neuron Response Suppression in Owl Monkey Primary Somatosensory Cortex When Stimuli Are Presented to Both Hands. J. Neurosci. 31, 3589–3601.

Rigotti, M., Barak, O., Warden, M.R., Wang, X.-J., Daw, N.D., Miller, E.K., and Fusi, S. (2013). The importance of mixed selectivity in complex cognitive tasks. Nature 497, 585–590.

Saal, H.P., Harvey, M.A., and Bensmaia, S.J. (2015a). Rate and timing of cortical responses driven by separate sensory channels. Elife 4.

Saal, H.P., Harvey, M.A., and Bensmaia, S.J. (2015b). Rate and timing of cortical responses driven by separate sensory channels. Elife 4.

Sinclair, R.J., and Burton, H. (1991). Neuronal activity in the primary somatosensory cortex in monkeys (Macaca mulatta) during active touch of textured surface gratings: responses to groove width, applied force, and velocity of motion. J. Neurophysiol. 66, 153–169.

Skedung, L., Arvidsson, M., Chung, J.Y., Stafford, C.M., Berglund, B., and Rutland, M.W. (2013). Feeling small: exploring the tactile perception limits. Sci. Rep. 3, 2617.

Sripati, A.P., Bensmaia, S.J., and Johnson, K.O. (2006b). A continuum mechanical model of mechanoreceptive afferent responses to indented spatial patterns. J. Neurophysiol. 95, 3852–3864.

Sripati, A.P., Yoshioka, T., Denchev, P., Hsiao, S.S., and Johnson, K.O. (2006a). Spatiotemporal receptive fields of peripheral afferents and cortical area 3b and 1 neurons in the primate somatosensory system. J. Neurosci. 26, 2101–2114.

Stringer, C., Pachitariu, M., Steinmetz, N., Reddy, C., Carandini, M., and Harris, K.D. (2018). Spontaneous behaviors drive multidimensional, brain-wide neural activity. BioRxiv 1–26.

Sur, M., Merzenich, M.M., and Kaas, J.H. (1980). Magnification, receptive-field area, and “hypercolumn” size in areas 3b and 1 of somatosensory cortex in owl monkeys. J. Neurophysiol. 44, 295–311.

Theriault, B.R., Reed, D.A., and Niekrasz, M.A. (2008). Reversible medetomidine/ketamine anesthesia in captive capuchin monkeys (Cebus apella). In Journal of Medical Primatology, pp. 74–81.

Tremblay, F., Ageranioti-Bélanger, S. a., and Chapman, C.E. (1996). Cortical mechanisms underlying tactile discrimination in the monkey. I. Role of primary somatosensory cortex in passive texture discrimination. J. Neurophysiol. 76.

Turner, E.C., Young, N.A., Reed, J.L., Collins, C.E., Flaherty, D.K., Gabi, M., and Kaas, J.H. (2016). Distributions of Cells and Neurons across the Cortical Sheet in Old World Macaques. Brain. Behav. Evol. 88, 1–13.

Weber, A.I., Saal, H.P., Lieber, J.D., Cheng, J.-W., Manfredi, L.R., Dammann, J.F., and Bensmaia, S.J. (2013). Spatial and temporal codes mediate the tactile perception of natural textures. Proc. Natl. Acad. Sci. U. S. A. 110, 17107–17112.

Yoshioka, T., Gibb, B., Dorsch, a K., Hsiao, S.S., and Johnson, K.O. (2001). Neural coding mechanisms underlying perceived roughness of finely textured surfaces. J. Neurosci. 21, 6905–6916.

